# ULDNA: Integrating Unsupervised Multi-Source Language Models with LSTM-Attention Network for Protein-DNA Binding Site Prediction

**DOI:** 10.1101/2023.05.30.542787

**Authors:** Yi-Heng Zhu, Dong-Jun Yu

## Abstract

Accurate identification of protein-DNA interactions is critical to understand the molecular mechanisms of proteins and design new drugs. We proposed a novel deeplearning method, ULDNA, to predict DNA-binding sites from protein sequences through a LSTM-attention architecture embedded with three unsupervised language models pretrained in multiple large-scale sequence databases. The method was systematically tested on 1287 proteins with DNA-binding site annotation from Protein Data Bank. Experimental results showed that ULDNA achieved a significant increase of the DNA-binding site prediction accuracy compared to the state-of-the-art approaches. Detailed data analyses showed that the major advantage of ULDNA lies in the utilization of three pre-trained transformer language models which can extract the complementary DNA-binding patterns buried in evolution diversity-based feature embeddings in residue-level. Meanwhile, the designed LSTM-attention network could further enhance the correlation between evolution diversity and protein-DNA interaction. These results demonstrated a new avenue for high-accuracy deep-learning DNA-binding site prediction that is applicable to large-scale protein-DNA binding annotation from sequence alone.

## 1. Introduction

Protein-DNA interactions play critical roles in various biological processes, including gene expression and regulation, DNA replication, repair, and recombination [1, 2]. The accurate identification of protein-DNA binding residues not only contributes to understanding the molecular mechanisms of proteins, but also has important practical significance for drug design [3]. Direct determination of DNA-binding sites via biochemical experiments, such as fast ChIP5 [4], X-ray crystallography [5], and Cryo-EM [6], is typically time-consuming and laborious, and often incomplete. As a result, numerous sequenced proteins have no available DNA-binding annotation to date. As of June 2023, for example, the UniProt database [7] harbored ∼246 million protein sequences, but only <0.1% of them were annotated with known DNA-binding site records using experimental evidence. To fill the gap between sequence and DNA-binding annotation, it is urgent to develop efficient computational methods for protein-DNA binding site prediction [8, 9].

Existing DNA-binding site prediction methods can be divided into two categories, i.e., template detection-based methods and machine learning-based methods [10]. In the early stage, template detection-based methods lead the trend of protein-DNA interaction prediction [11, 12]. Specifically, these methods identify DNA-binding sites through detecting the templates that have similar sequence or structure to the query. For examples, S-SITE [13] identifies sequence templates using PSI-BLAST alignment [14], while PreDNA [15] and DBD-Hunter [16] search templates through structure alignment. There exist other elegant predictors, including PreDs [17], DBD-Threader [18], DR_bind [19], and Morozov’s method [20].

Template-based approaches have a common drawback: the accuracy of these methods is contingent upon the availability of templates with readily identifiable DNA-binding site annotation. To eliminate such dependence, machine learning-based methods have emerged to extract hand-crafted features from sequences and structures (e.g., position-specific scoring matrix [21] and solvent accessible surface area [22]), which can then be used by machine learning approaches (e.g., support vector machine [23] and random forest [24]) to implement DNA-binding site prediction, with typical examples including DNAPred [10], TargetDNA [25], MetaDBSite [26], and TargetS [27].

Despite the potential advantage, the prediction accuracy of many early machine learning-based methods was not satisfactory. One of the major reasons is due to the lack of informative feature representation methods, as most of the approaches are based on simple feature representations, such as amino acid coding, physiochemical properties, and evolution conservation, which cannot fully extract the complex pattern of protein-DNA interaction [28, 29]. To partly overcome this barrier, several methods, e.g., Guan’s method [30], PredDBR [31], iProDNA-CapsNet [32], and GraphBind [33], utilize deep learning technology to predict DNA-binding sites. Compared to traditional machine learning approaches, one advantage of deep learning technologies is that they could extract more discriminative feature embeddings from sequences and structures through designing complex neural networks. Nevertheless, the performance of deep learning methods is often hampered by the limitation of experimental annotation data consisting of only ∼4000 protein-DNA complexes from Protein Data Bank (PDB) [34]. The insufficient experimental data significantly limit the effectiveness of training the deep neural network models.

To alleviate the issue caused by the lack of annotated data, a promising approach is to utilize protein language models pre-trained through deep-learning networks on large-scale sequence databases without DNA-binding annotations. Due to the extensive sequence training and learning, important inter-residue correlation patterns, which are critical for protein-DNA interaction, can be extracted through the language models and utilized for feature embedding. Recently, several protein language models, such as TAPE [35], SeqVec [36], and Bepler’s approach [37], have been emerged, often through supervised learners such as convolutional neural networks (CNNs) [38], in protein structure and function prediction tasks, with examples including the predictions of contact map [39], molecular function [40], mutation and stability [35], and GO transferals [41].

In this work, we proposed a new deep learning model, ULDNA, for high accuracy protein-DNA binding site prediction by the integration of the unsupervised protein language models from multiple information sources with the designed LSTM-attention network. Specifically, we utilize three recently proposed language models (i.e., ESM2 [42], ProtTrans [43], and ESM-MSA [44]), separately pre-trained on different large-scale sequence databases, to extract the complementary evolution diversity-based feature embeddings, which are highly associated with protein-DNA interaction. Then, a LSTM-attention network is designed to train DNA-binding site models from multi-source feature embeddings through enhancing the correlation between evolution diversity and DNA-binding pattern. ULDNA has been systematically tested on five protein-DNA binding site datasets, where the results demonstrated significant advantage on accurate DNA-binding site prediction over the current state-of-the-art of the field. The standalone package and an online server of ULDNA are made freely available through URL http://csbio.njust.edu.cn/bioinf/uldna/.

## 2. Materials and methods

### 2.1 Benchmark datasets

The proposed methods were evaluated by five protein-DNA binding site datasets, including PDNA-543 [25], PDNA-41 [25], PDNA-335 [27], PDNA-52 [27], and PDNA-316 [26], from previous works.

PDNA-543 and PDNA-41 separately consist of 543 and 41 DNA-binding protein chains, which were deposited in the PDB before and after October 10, 2014, respectively. Here, a sequence identity cut-off 30% has been used to filter out the redundant proteins within each dataset and between different datasets using the CD-HIT program [45].

PDNA-335 and PDNA-52 contain 335 and 52 DNA-binding chains, respectively, which are released in PDB before and after March 10, 2010. The sequence identity within each dataset and between different datasets is reduced to 40% through PISCES software [46].

PDNA-316 collects 316 DNA-binding chains before 2011, where the maximal pairwise sequence identity of proteins is culled to 30% using CD-HIT [45]. The detailed statistical summary of five datasets is presented in Table 1.

**Table 1.** Statistical summary of five protein-DNA binding site datasets.

| Dataset | No. of Sequences | No. of DNA-binding residues | No. of No-DNA-binding residues |
| --- | --- | --- | --- |
| PDNA-543 | 543 | 9,549 | 134,995 |
| PDNA-41 | 41 | 734 | 14,021 |
| PDNA-335 | 335 | 6,461 | 71,320 |
| PDNA-52 | 52 | 973 | 16,225 |
| PDNA-316 | 316 | 5,609 | 67,109 |

### 2.2 The framework of ULDNA

ULDNA is a deep learning-based method for protein-DNA binding site prediction, with input being a query amino acid sequence and output including confidence scores of belonging DNA-binding sites. As shown in Figure 1, ULDNA consists of two procedures, including feature embedding extraction using multi-source language models and DNA-binding site prediction using LSTM-attention network.

**Figure 1.**
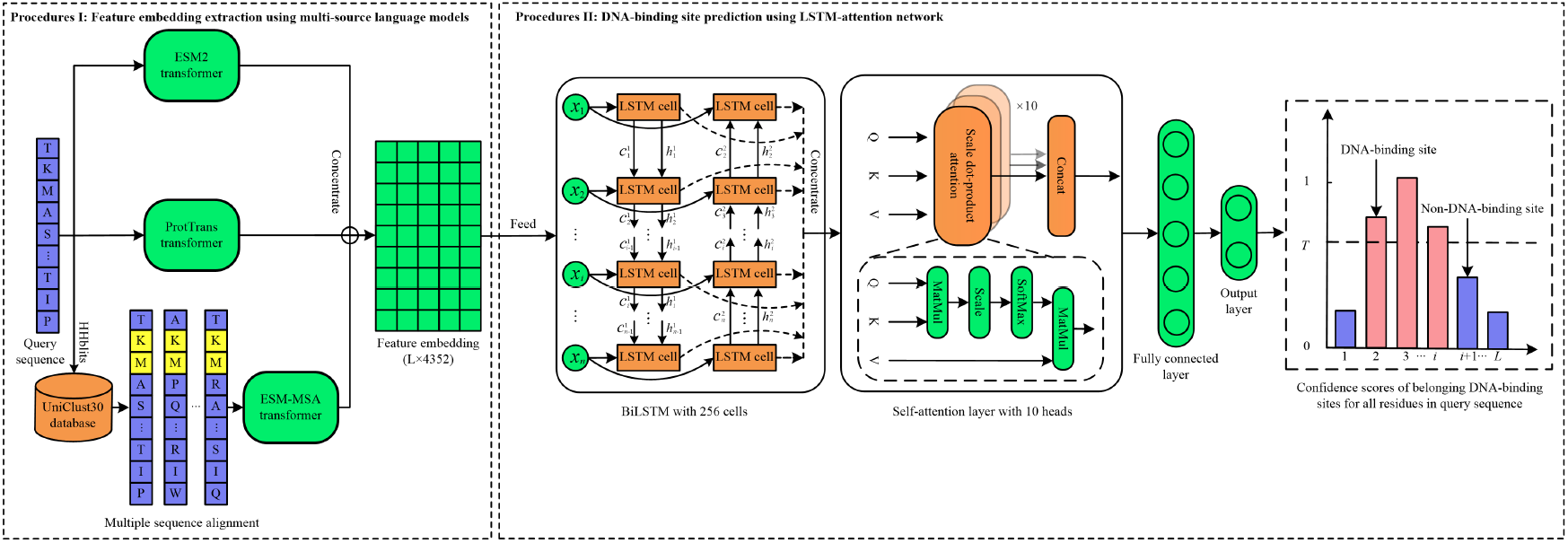
The workflow of ULDNA **Procedure I: Feature embedding extraction using multi-source language models**. The input sequence is fed to ESM2 [42] and ProtTrans [43] transformers to output two feature embedding matrices with the scales of *L* × 2560 and *L* × 1024, respectively; Meanwhile, we search the multiple sequence alignment (MSA) of the input sequence from UniClust30 database [47], which is further fed to ESM-MSA transformer [44] to generate another feature embedding matrix with the scale of *L* × 768, where *L* is the length of input sequence, 2560, 1024 and 768 are preset hyper-parameters in transformer models. ESM2, ProtTrans, and ESM-MSA are both unsupervised attention networks with 36, 24, and 12 layers, respectively, and separately trained on Uniref50 [48], UniClust30 & Uniref50, and BFD (Big Fantastic Database) [49] & Uniref50, respectively, where “&” means that two databases are both used to train a transformer. Each transformer has learnt abundant evolution knowledge from millions of sequences and could encode the input sequence (or MSA) as a feature embedding matrix with evolution diversity. Considering that the evolution knowledge from multiple database sources could be complementary, we concentrate the above-mentioned three feature embedding matrices from different transformer models as a combination embedding matrix with the scale of *L* × 4352. **Procedure II: DNA-binding site prediction using LSTM-attention network**. The concentrated feature embedding is fed to a designed LSTM-attention network to generate a score vector with *L* dimensions, indicating the confidence scores of belonging DNA-binding sites for all residues in query sequence. In LSTM-attention network, a BiLSTM layer and a self-attention layer are combined to enhance the correlation between evolution diversity and DNA-binding in residue-level to improve DNA-binding prediction.

### 2.3 Unsupervised protein language models

The architecture of ESM2 transformer [42] is illustrated in Figure S1, with input and output being a query amino acid sequence and an evolution diversity-based feature embedding matrix, respectively. ESM2 includes 36 attention layers, each of which consists of 20 attention heads and a feed-forward network (FFN). In each attention head, the scale dot-product attention is performed to learn the evolution correlation between amino acids in the query sequence from an individual view. Then, the FFN fuses the evolution knowledge from all attention heads to capture the evolution diversity for the entire sequence. The ESM2 model with 3 billion parameters was trained on over 60 million proteins from UniRef50 database, as carefully described in Text S1.

ProtTrans transformer [43] shares the similar architecture to ESM2, with including 24 layers, each of which consists of 32 attention heads. The ProtTrans model with 3 billion parameters was trained on over 45 million proteins from BFD and UniRef50 databases.

ESM-MSA transformer [44] is designed to extract the co-evolution-based feature embedding matrix for a MSA, as shown in Figure S2. ESM-MSA consists of 12 attention blocks, each of which contains a row-attention layer and a column-attention layer which separately learn the co-evolution correlation between amino acids in sequence-level and position-level. The ESM-MSA model with 100 million parameters was trained on over 26 million MSAs from Unclust30 and UniRef50 databases, with details in Text S2.

### 2.4 LSTM-attention network

As shown in Figure 1, the designed LSTM-attention network consists of a BiLSTM layer, a self-attention layer, a fully connected layer, and an output layer.

The BiLSTM layer includes a forward LSTM and a backward LSTM, which have the same architecture consisting of 256 cells with reverse propagation directions. Each LSTM cell is mainly composed of two states (i.e., cell state *c* and hidden state *h*) and three gates (i.e., forget gate *f*, input gate *i*, and output gate *o*). Cell state and hidden state are separately used to store and output the signals at the current time-step. Forget gate, input gate, and output gate are used to control the ratios of incorporating history signal, inputting current signal, and outputting updated signal, respectively. Specifically, at time-step *t* (*t* ≤ *L, L* is the length of input sequence), the above-mentioned states and gates are computed as follows:

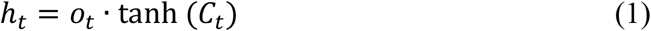

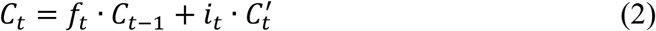

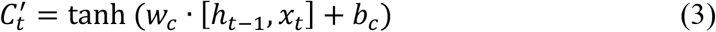

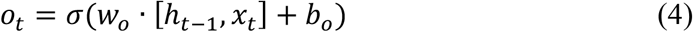

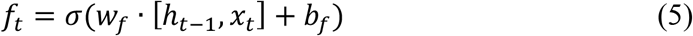

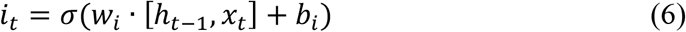

where *C*_*t*‒1_ and *h*_*t*‒1_ are cell state and hidden state, respectively, at the time-step *t* − 1, *x*_*t*_ is the input vector at the time-step *t* (i.e., the feature embedding vector with 4352 dimensions of the *t* -th residue in the query sequence for DNA-binding prediction), *w*_∗_ is the weight vector, *b*_∗_ is the bias, [,] is concentration operation between two vectors, and σ is the Sigmoid function. The output of BiLSTM layer is represented as a *L* × 512 matrix through concentrating the hidden states in all LSTM cells at all time-steps.

The self-attention layer consists of 10 attention heads, each of which performs the scale dot-product attention as follows:

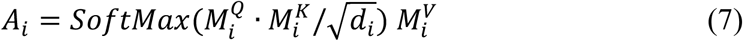

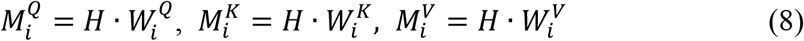

where *H* is the output matrix by the BiLSTM layer, *A*_*i*_ is a *L* × 64 attention matrix in the *i*-th attention head, 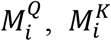, and 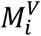 are Query, Key, and Value matrices, respectively, 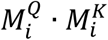 is a *L* × *L* weight matrix measuring the position-correlation for each amino acid pair in the query sequence, and *d*_*i*_ is the scale parameter.

The attention matrices in all of 10 heads are concentrated and then fed to a fully connected layer with 1024 neurons, followed by an output layer with 1 neuron:

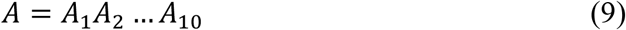

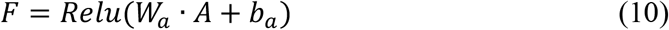

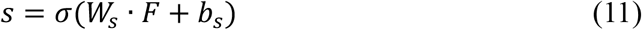

where *s* is a score vector with *L* dimensions, indicating the confidence scores of belonging DNA-binding sites for all residues for the query sequence.

The cross-entropy loss [50] is used as the training loss:

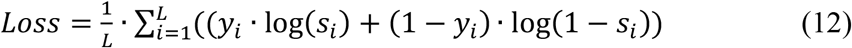

where *s*_*i*_ is the confidence score of belonging DNA-binding site for the *i*-th residue in the query sequence; *y*_*i*_ = 1, if the *i* -th residue is a DNA-binding site in the experimental annotation; otherwise, *y*_*i*_ = 0. We minimize the loss function to optimize ULDNA pipeline using Adam optimization algorithm [51].

### 2.5 Evaluation indices

Four indices, i.e., Sensitivity (Sen), Specificity (Spe), Accuracy (Acc), and Mathew’s Correlation Coefficient (MCC), are utilized to evaluate the proposed methods:

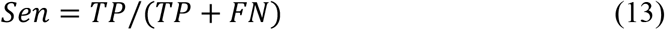

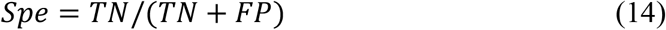

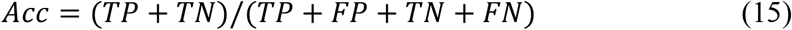

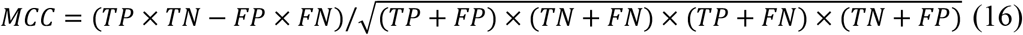

where TP, TN, FP, and FN represent true positives, true negatives, false positives, and false negatives, respectively.

Because the above four indices are threshold-dependent, it is critical to select an appropriate threshold for fair comparisons between various predictors. In this work, we select the threshold which maximizes the value of MCC over ten-fold cross-validation. Moreover, a threshold-independent index, i.e., area under the receiver operating characteristic curve (AUROC), is used to evaluate the overall prediction performances of predictors.

## 3. Results and discussions

### 3.1 Comparison with existing DNA-binding site predictors

To demonstrate the strong performance of the proposed ULDNA, we compared it with 12 start-of-the-art DNA-binding site predictors, including BindN [52], ProteDNA [53], BindN+ [54], MetaDBSite [26], DP-Bind [55], DNABind [56], TargetDNA [25], iProDNA-CapsNet [32], DNAPred [10], Guan’s method [30], COACH [13], and PredDBR [31], on PDNA-41 test dataset under independent validation, as shown in Table 1. It could be found that ULDNA achieves the highest MCC values among all of 13 competing methods. Compared to the second-best performer, i.e., PredDBR (a recently proposed deep learning model), ULDNA gains 43.9% average improvement of MCC values under three thresholds, respectively. More importantly, four evaluation metrics of ULDNA are both higher than those of PredDBR under *Sen* ≈ *Spe* and *Spe* ≈ 0.95. Meanwhile, a similar trend but with more significant distinctions can be observed in comparison with other predictors. Taking DNAPred as an example, ULDNA shares 9.3%, 18.6%, 17.6%, 76.2%, and 9.0% improvements for Sen, Spe, Acc, MCC, and AUROC values, respectively, under *Sen* ≈ *Spe*. It cannot escape from our notice that ProteDNA obtains the highest Spe value (0.998) but with the lowest Sen (0.048). The underlying reason is that ProteDNA predict too many false negatives.

**Table 1.** Performance comparisons between ULDNA and 12 competing predictors on PDNA-41 under independent validation.

| Method | Sen | Spe | Acc | MCC | AUROC |
| --- | --- | --- | --- | --- | --- |
| BindN <sup>a</sup> | 0.456 | 0.809 | 0.792 | 0.143 | - |
| ProteDNA <sup>a</sup> | 0.048 | **0.998** | 0.951 | 0.160 | - |
| BindN+ (*Spe* ≈ 0.95) <sup>a</sup> | 0.241 | 0.951 | 0.916 | 0.178 | - |
| BindN+ (*Spe* ≈ 0.85) <sup>a</sup> | 0.508 | 0.854 | 0.837 | 0.213 | - |
| MetaDBSite <sup>a</sup> | 0.342 | 0.934 | 0.904 | 0.221 | - |
| DP-Bind <sup>a</sup> | 0.617 | 0.824 | 0.814 | 0.241 | - |
| DNABind <sup>a</sup> | 0.702 | 0.803 | 0.798 | 0.264 | - |
| TargetDNA (*Sen* ≈ *Spe*) <sup>a</sup> | 0.602 | 0.858 | 0.845 | 0.269 | - |
| TargetDNA (*Spe* ≈ 0.95) <sup>a</sup> | 0.455 | 0.933 | 0.909 | 0.300 | - |
| iProDNA-CapsNet (*Sen* ≈ *Spe*) <sup>b</sup> | 0.754 | 0.753 | 0.753 | 0.245 | - |
| iProDNA-CapsNet (*Spe* ≈ 0.95) <sup>b</sup> | 0.422 | 0.949 | 0.924 | 0.315 | - |
| DNAPred (*Sen* ≈ *Spe*) <sup>c</sup> | 0.761 | 0.767 | 0.761 | 0.260 | 0.858 |
| DNAPred (*Spe* ≈ 0.95) <sup>c</sup> | 0.447 | 0.949 | 0.924 | 0.337 | 0.858 |
| Guan’s method <sup>d</sup> | 0.476 | 0.964 | 0.949 | 0.357 | - |
| COACH <sup>e</sup> | 0.462 | 0.951 | 0.927 | 0.352 | - |
| PredDBR (*Sen* ≈ *Spe*) <sup>e</sup> | 0.764 | 0.758 | 0.758 | 0.264 | - |
| PredDBR (*Spe* ≈ 0.95) <sup>e</sup> | 0.431 | 0.958 | 0.931 | 0.351 | - |
| PredDBR (threshold = 0.5) <sup>e</sup> | 0.391 | 0.968 | 0.939 | 0.359 | - |
| ULDNA (*Sen* ≈ *Spe*) | **0.824** | 0.899 | 0.895 | 0.458 | **0.935** |
| ULDNA (*Spe* ≈ 0.95) | 0.556 | 0.970 | 0.950 | **0.499** | **0.935** |
| ULDNA (threshold = 0.5) | 0.271 | 0.994 | **0.958** | 0.417 | **0.935** |

<sup>a</sup> Results excerpted from TargetDNA [25]; <sup>b</sup> Results excerpted from iProDNA-CapsNet [32]; <sup>c</sup> Results excerpted from DNAPred [10]; <sup>d</sup> Results excerpted from Guan et al [30]; <sup>e</sup> Results excerpted from PredDBR [31]; “*Sen* ≈ *Spe*” and “*Spe* ≈ 0.95” mean the thresholds that make *Sen* ≈ *Spe* and “*Spe* ≈ 0.95”, respectively, on PDNA-543 training dataset over ten-fold cross-validation. Bold fonts highlight the best performer in each evaluation metric.

Table 2 summarizes the performance comparison among DNABR [29], MetaDBSite [26], TargetS [27], DNAPred [10], COACH [13], PredDBR [31], and ULDNA on PDNA-52 test dataset under independent validation, where ULDNA achieves the highest MCC value among all control methods. Specifically, the improvements of MCC values between ULDNA and other 6 predictors range from 14.6% to 179.5%.

**Table 2.**
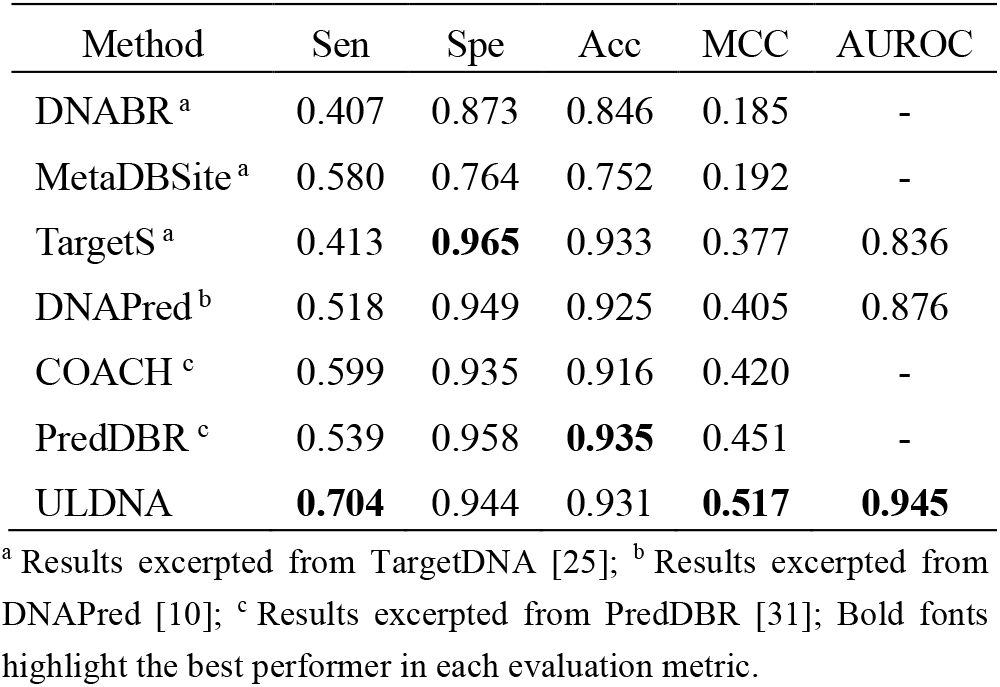
Performance comparisons between ULDNA and 6 competing predictors on PDNA-52 under independent validation.

| Method | Sen | Spe | Acc | MCC | AUROC |
| --- | --- | --- | --- | --- | --- |
| DNABR <sup>a</sup> | 0.407 | 0.873 | 0.846 | 0.185 | - |
| MetaDBSite <sup>a</sup> | 0.580 | 0.764 | 0.752 | 0.192 | - |
| TargetS <sup>a</sup> | 0.413 | **0.965** | 0.933 | 0.377 | 0.836 |
| DNAPred <sup>b</sup> | 0.518 | 0.949 | 0.925 | 0.405 | 0.876 |
| COACH <sup>c</sup> | 0.599 | 0.935 | 0.916 | 0.420 | - |
| PredDBR <sup>c</sup> | 0.539 | 0.958 | **0.935** | 0.451 | - |
| ULDNA | **0.704** | 0.944 | 0.931 | **0.517** | **0.945** |

<sup>a</sup> Results excerpted from TargetDNA [25]; <sup>b</sup> Results excerpted from DNAPred [10]; <sup>c</sup> Results excerpted from PredDBR [31]; Bold fonts highlight the best performer in each evaluation metric.

We further compare our method with all the above mentioned methods as well as other 4 competing methods, including EC-RUS [57], DBS-PRED [58], DISIS [59] and BindN-rf [28], on three training datasets (i.e., PDNA-543, PDNA-335, and PDNA-316) over cross-validation, as listed in Tables S1, S2, and S3. Again, the proposed ULDNA outperforms all other methods.

### 3.2 Contribution analysis of different protein language models

To analyze the contributions of three protein language models (i.e., ESM2, ProtTrans, and ESM-MSA) in DNA-binding site prediction, we further benchmarked the designed LSTM-attention network with seven different feature embeddings, respectively, including three individual embeddings from ESM2, ProtTrans, and ESM-MSA, and four combination embeddings from ProtTrans + ESM-MSA (PE), ESM2 + ESM-MSA (EE), ESM2 + ProtTrans (EP), and ESM2 + ProtTrans + ESM-MSA (EPE), where “+” means the individual feature embeddings from different language models are concentrated as a combination embeddings. Figure 1 illustrates the performance of seven feature embeddings on three training datasets (i.e., PDNA-543, PDNA-335, and PDNA-316) under cross-validation and two test datasets (i.e., PDNA-41 and PDNA-52) under independent validation.

It could be found that EPE achieves the best performance among seven feature embeddings. From the view of MCC values, EPE separately gains 5.8%, 8.8%, 13.1%, 3.2%, 2.4%, and 2.0% average improvements on five datasets in comparison with ESM2, ProtTrans, ESM-MSA, PE, EE, and EP, respectively. With respect to AUROC values, EPE occupies the top-1 positions on four out of five datasets. Moreover, ESM2 shows the highest MCC and AUROC values among three individual embeddings; meanwhile, the largest increase is caused by adding ESM2 to PE on each dataset.

These data demonstrate the following two conclusions. First, three language models pretrained on different large-scale sequence databases are complementary to improve DNA-binding site prediction. Second, ESM2 made the most important contribution among three language models.

**Figure 1.**
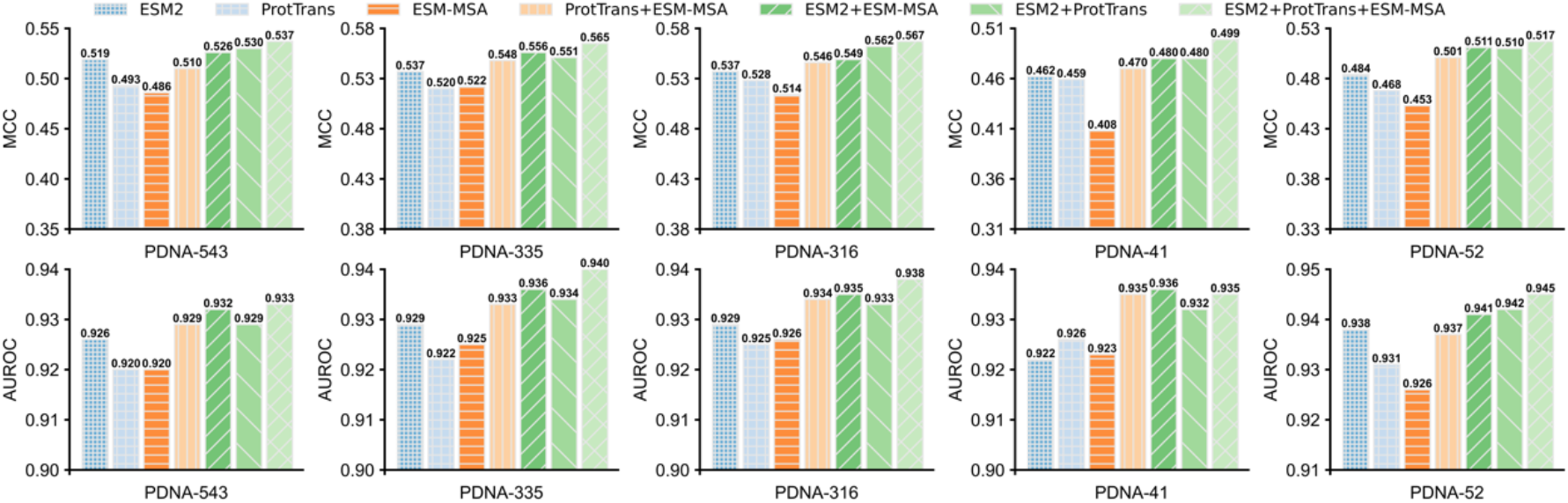
The MCC and AUROC values of seven feature embeddings on five benchmark datasets.

### 3.3 Ablation study

To analyze the contributions of algorithmic innovations in ULDNA to its improved performance, we design an ablation study, in which we start from a baseline model (M0) and incrementally add algorithmic components of ULDNA to build two advanced models (M1 and M2, with M2 = ULDNA). The pipelines of the three models are designed as follows (see Figure S1 for the architectures):

**M0**: Model is trained on BiLSTM consisting of 256 cells with a one-hot coding matrix [60] extracted from the input sequence, followed by a fully connected network, in which an output layer with SoftMax function [61] is used to generate the confidence scores of belonging DNA-binding sites for all residues in the input sequence. In the training stage, the cross-entropy loss [62] is used as the loss function, as described in Eq 9.

**M1**: We replace the one-hot coding matrix by a combination feature embedding matrix concentrated by three individual embeddings from ESM2, ProtTrans, and ESM-MSA. This combination embedding is further fed to the BiLSTM architecture used in M0 to output the confidence scores of belonging DNA-binding sites.

**M2 (M2=ULDNA)**: We add a self-attention layer consisting of 10 attention heads and a feed-forward network in the BiLSTM used in M1.

Figure 2 summarizes the performance of three ablation models on three training datasets under cross-validation and two test datasets under independent validation, where we run each model for 10 times and then used the average of all predictions as the final-result. Compared with M0, M1 achieves a significant gain with the average MCC and AUROC values increased by 148.4% and 23.4%, respectively, on five datasets, demonstrating that the protein language models are critical to improve DNA-binding site prediction of the ULDNA pipeline. After adding the self-attention layer in M1, the corresponding MCC values are increased on average by 1.3% on five datasets. The AUROC values of M2 cannot be further improved and even be slightly degraded on PDNA-543 and PDNA-41 in comparison with M1, but the corresponding values are sustainably increased on other three datasets. These observations indicate that the additional self-attention layer is helpful for enhancing the overall performance of function prediction, although less significant than the protein language models.

**Figure 2.**
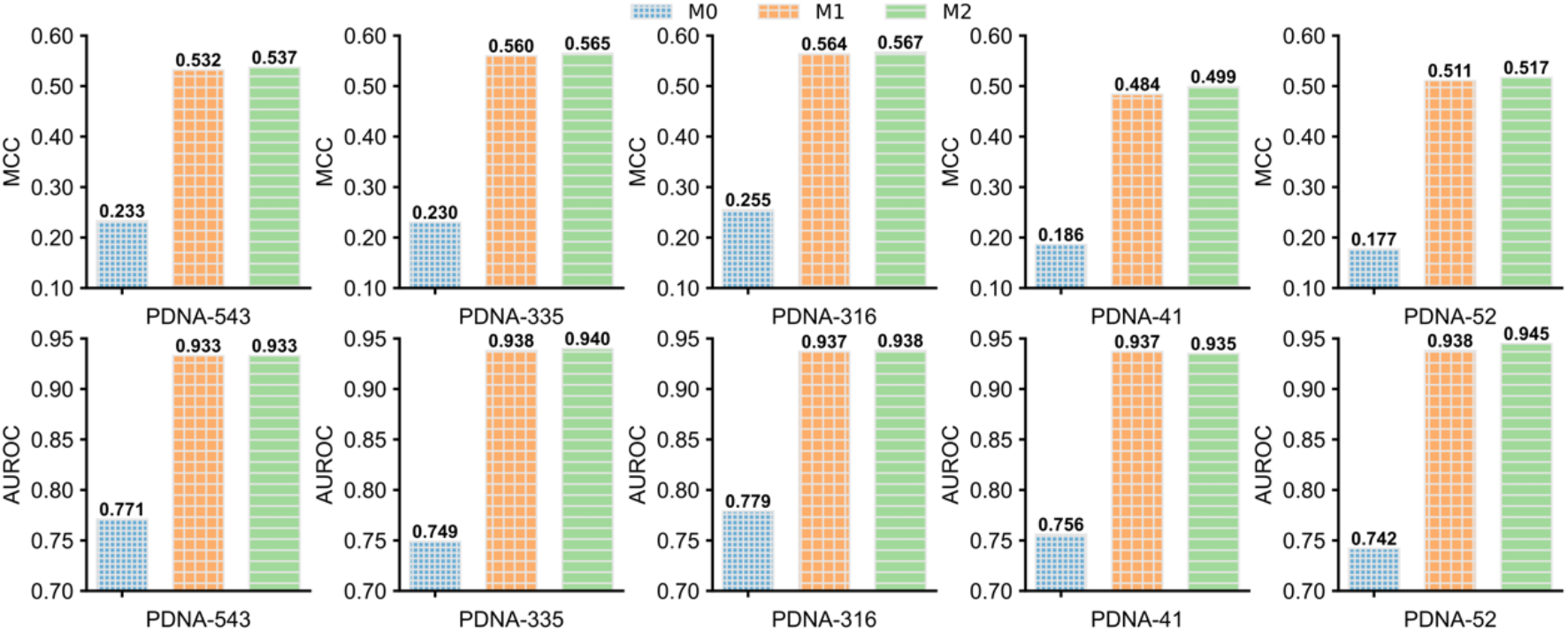
The MCC and AUROC values of three ablation models on five benchmark datasets.

### 3.4 Case study

To further examine the effects of different DNA-binding site prediction methods, we selected two proteins (2MXF_A and 3ZQL_A) from our test datasets as illustrations. For each protein, we used four in-house methods (denoted as LA-ESM2, LA-ProtTrans, LA-EMS-MSA-1b, and ULDNA) and a competing method (PredDBR [31]) to predict the corresponding DNA-binding sites. Four in-house methods use the same LSTM-attention network with different feature embeddings from ESM2, ProtTrans, ESM-MSA, and ESM2+ProtTrans+ESM-MSA, respectively, where “+” means the individual feature embeddings from different language models are concentrated as a combination embedding. Table 3 summarizes the modeling results of two proteins for five DNA-binding site prediction methods, where the corresponding visualization results are illustrated in Figure 3. In addition, the predicted and native DNA-binding sites of two proteins by five methods are listed in Table S4.

**Table 3.**
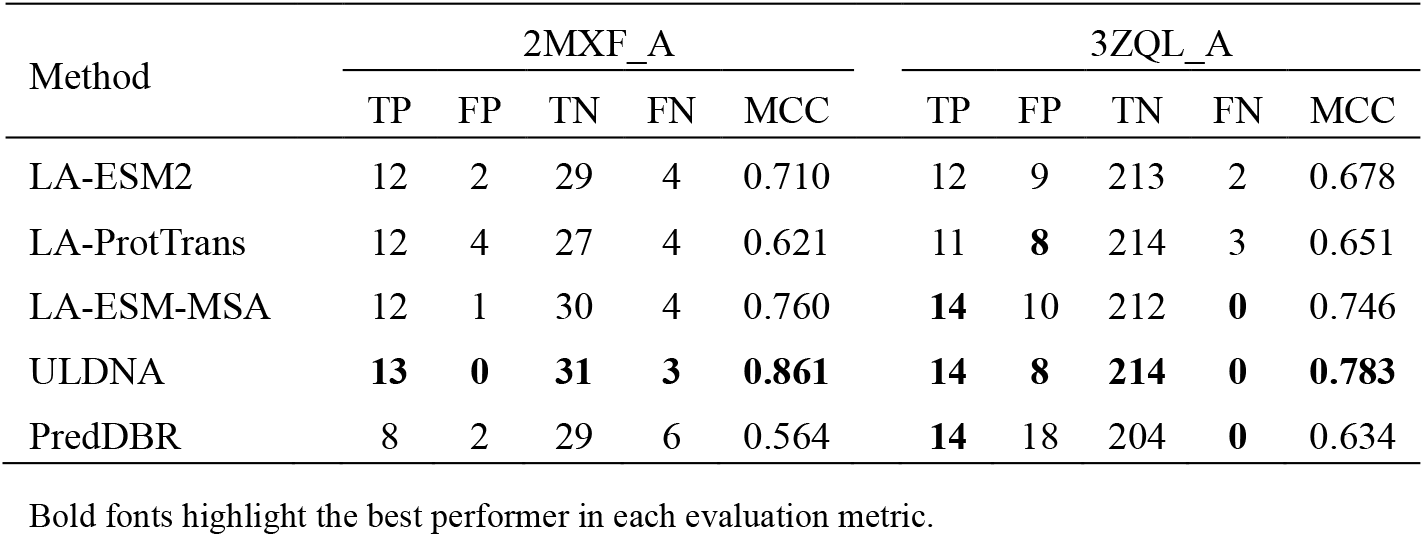
The modeling results of five DNA-binding site prediction methods on two representative examples 2MXF_A 3ZQL_A

**Figure 3.**
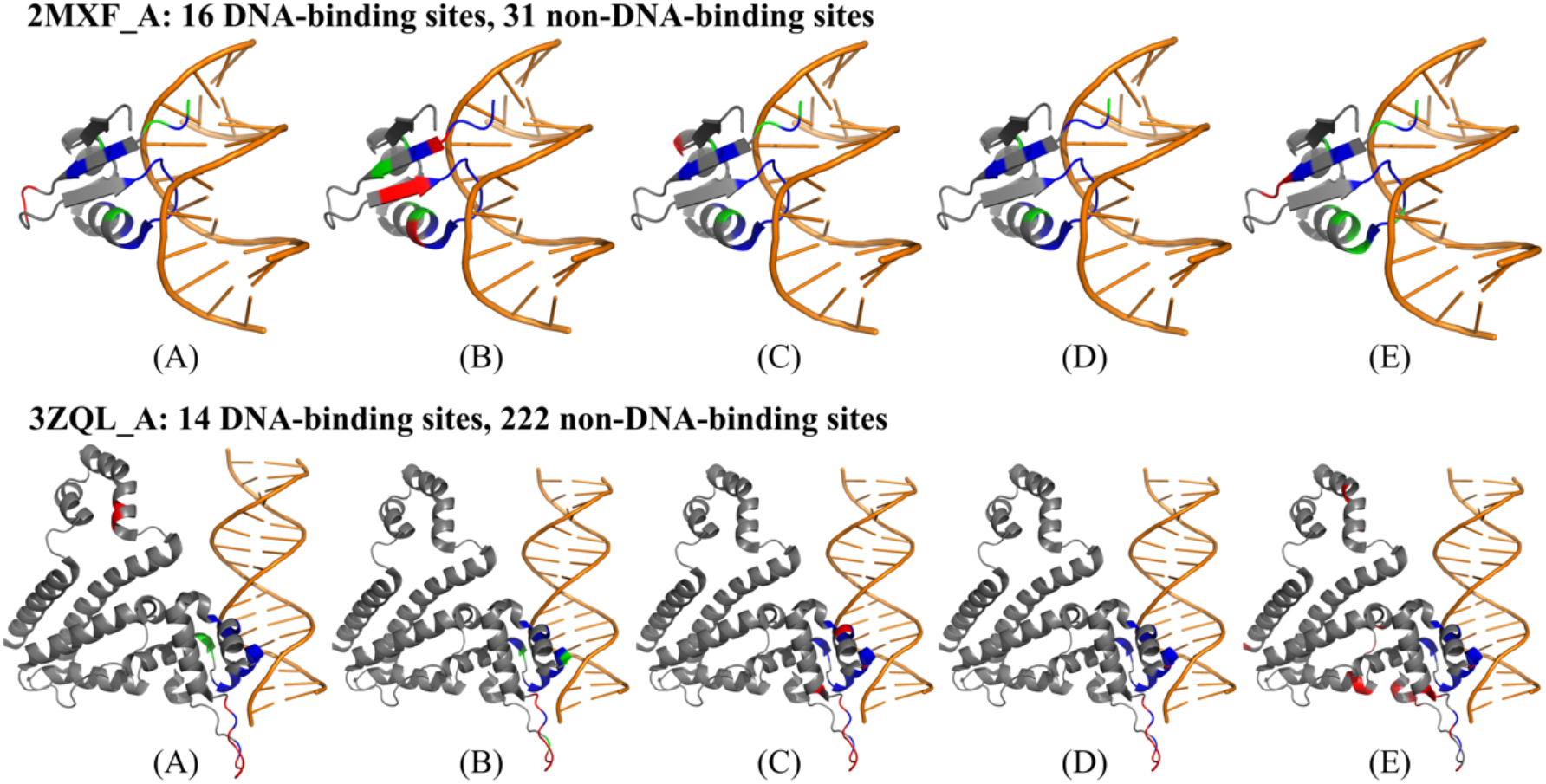
Visualization of prediction results for two proteins using five DNA-binding site prediction models: (A) LA-ESM2, (B) LA-ProtTrans, (C) LA-ESM-MSA, (D) ULDNA, (E) PredDBR. The color scheme is used as follows: DNA in orange, true positives in blue, false positives in red, false negatives in green. The pictures are made with PyMOL [63].

Several interesting observations can be made from the data. First, the protein language models are critical to improve DNA-binding site prediction. Specifically, each of four in-house methods with pre-trained protein language models shows the higher MCC values than the competing PredDBR without language models on two proteins.

Taking ULDNA as an example, it gains 52.7% and 23.5% improvements of MCC values on 2MXF and 3ZQL_A, respectively, in comparison with PredDBR.

Second, the combination of complementary protein language models can further increase the accuracy of ULDNA. In 2MXF_A, three in-house methods (i.e., LA-ESM2, LA-ProtTrans, and LA-ESM-MSA) with different language models hit 14 true positives in total, which is more than that by each individual method, indicating that three language models (i.e., ESM2, ProtTrans, and ESM-MSA) derive complementary information from different database sources. Meanwhile, the false positives predicted by one in-house method can be corrected by other two methods. For example, LA-ESM2 generates two false positives (10P and 11H), which are correctly predicted as non-DNA-binding sites by LA-ProtTrans and LA-ESM-MSA. As a result, by taking the combination of three language models, ULDNA gains the most-true positives without false positives among all methods. Sometimes, one in-house method can cover all true positives predicted by other methods. For example, for 3ZQL_A, all true positives of LA-ESM2 and LA-ProtTrans are covered by LA-ESM-MSA. Even in this case, the accuracy of final ULDNA is still improved by including the less accurate methods to reduce false positives.

## 4. Conclusions

We developed a new deep learning-based method, named ULDNA, to predict DNA-binding sites from the primary protein sequences. The algorithm was built on transformer embedding and LSTM-attention decoding. The large-scale tests on five protein-DNA binding site datasets demonstrated that ULDNA consistently outperforms other state-of-the-art approaches in the accuracy of DNA-binding site prediction. The improvement of ULDNA can be attributed to several advancements. First and most importantly, three transformers can effectively extract complementary evolution diversity-based feature embeddings for the input sequence from different database sources. Second, the designed LSTM-attention network enhances the correlation between evolution diversity and protein-DNA interaction pattern to improve prediction accuracy.

Despite the encouraging performance, there is still considerable room for further improvements. First, the serial feature concentration strategy, currently used in ULDNA, cannot perfectly deal with the redundant information among the feature embeddings from different transformers, where a more advanced feature fusion approaches may alleviate the negative impact caused by information redundancy in the future. Second, with the development of protein structure prediction models (e.g., AlphaFold2 [64] and ESMFold [42]), the predicted structures will have the huge potential to improve DNA-binding site prediction. Studies along these lines are under progress.

## Supporting Texts

### Text S1. The procedures for ESM2 transformer

#### A. Masking

For an input sequence, the masking strategy [1] is performed on the corresponding tokens (i.e., amino acids). Specifically, we randomly sample 15% tokens, each of which is changed as a special “masking” token with 80% probability, a randomly chosen alternate amino acid with 10% probability, and the original input token (i.e., no change) with 10% probability.

#### B. One-hot encoding

The masked sequence is represented as a *L* × 28 matrix using one-hot encoding [2], where 28 is the types of tokens, including 20 common amino acids, 6 non-common amino acids (B, J, O, U, X and Z), 1 gap token, and 1 “masking” token.

#### C. Embedding with position information

The one-hot coding matrix *X* of the masked sequence is multiplied by an embedding weight matrix *W*_*t*_ to generate an embedding matrix *H*_*t*_:

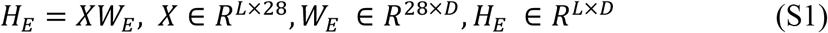

where *L* is the length of the masked sequence, 28 is the types of tokens in the masked sequence, and *D* is the embedding dimension.

Then, the position embedding strategy is used to record the position of each token in the masked sequence to generate a position embedding matrix *H*_*P*_:

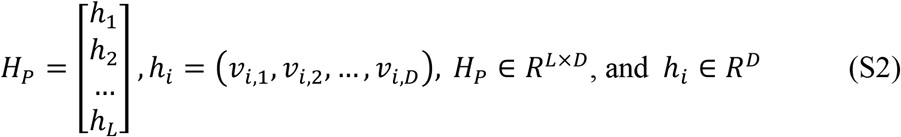

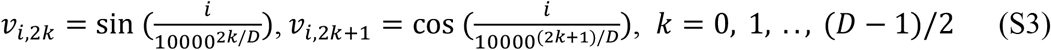

where *h*_*i*_ is the embedding vector for the *i*-th position in the masked sequence.

Finally, two embedding matrices are added as a combination embedding matrix *H*_1_:

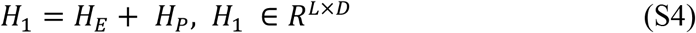

#### D. Self-attention

The embedding matrix *H*_*i*_ is fed to a self-attention block with *n* layers, each of which consists of *m* attention heads, a linear unit, and a feed-forward network (FFN). In each attention head, the scale dot-product attention is performed as follows:

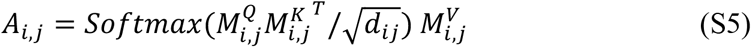

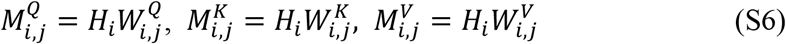

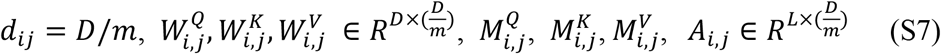

where *A*_*i,j*_ is the attention matrix in the (*i* -th layer, *j*-th head) and measures the evolution correlation for each amino acid pair in the sequence, 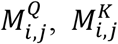, and 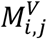 are Query, Key, and Value matrices in the (*i*-th layer, *j*-th head), *H*_*i*_ is the input matrix in the *i*-th layer, 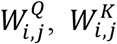, and 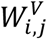 are weight matrices, and *d*_*ij*_ is the scale parameter.

The outputs of all attention heads in *i*-th layer are concatenated as a new matrix *A*_*i*_, which is further fed to a linear unit to output the matrix *U*_*i*_ :

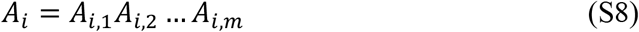

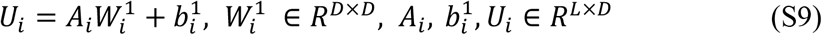

where 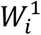 and 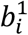 are the weight matrix and bias, respectively, in the linear unit.

#### E. Feed-forward network with shortcut connections

The *U*_*i*_ is added by *H*_*i*_ to generate a new matrix *F*_*i*_, which is further fed to the FFN to output the matrix *T*_*i*_:

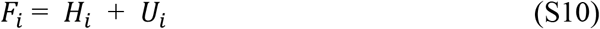

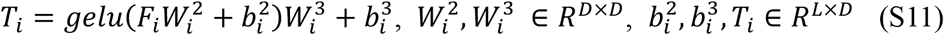

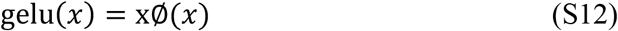

where 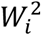 and 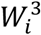 are weight matrices in the FFN, 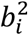 and 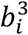 are bias in the FFN, and ∅(*x*) is the integral of Gaussian Distribution for *x*

The *F*_*i*_ is added by *T*_*i*_ as the output the *i*-th attention layer:

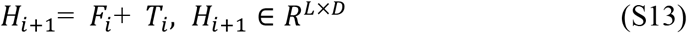

where *H*_*i*+1_ is the evolution diversity-based embedding matrix in *i*-th attention layer.

The output of the last attention layer is fed to a fully connected layer with SoftMax function to generate a *L* × 28 probability matrix:

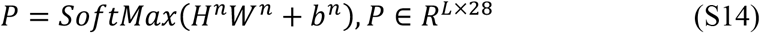

where the (*l*-th, *c*-th) value in *P* indicates the probability that the *l*-th token in the masked sequence is predicted as the *c*-th type of amino acid, *W*^*O*^ and *b*^*O*^ are weight matrix and bias, respectively.

#### F. Loss function

The loss function is designed as a negative log likelihood function between inputted one-hot and outputted probability matrices, to ensure that the prediction model correctly predicts the true amino acids in the masked position as much as possible:

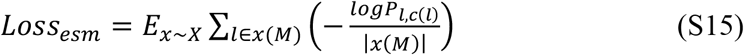

where *x* is a sequence in training protein set *X, x*(*M*) is a set of masking position in *x*, |*x*(*M*)| is the number of elements in *x*(*M*), *c*(*l*) is the type index of amino acid for the *l*-th token in *x* before masking, and –*logP*_l,c(l)_ is negative log likelihood of the true amino acid *x*_l_ under condition of masking.

The ESM2 transformer is optimized by minimizing the loss function via Adam optimization algorithm [3]. Then, the output of last attention layer is represented as a *L* × *D* matrix, as the evolution diversity-based embedding for DNA-binding site prediction, where *D* is the number of neurons of FFN. The current ESM2 model with 3 billion parameters was trained over 60 million proteins from UniRef50 database and can be freely download at https://github.com/facebookresearch/esm, where *n* = 36, *m* = 20, and *D* = 2560.

### Text S2. The procedures for ESM-MSA transformer

#### A. Masking

For an input multiple sequence alignment (MSA), the masking strategy is performed. Specifically, for each individual sequence in MSA, we randomly sample 15% tokens (amino acids), each of which is changed as a special “masking” token with 80% probability, a randomly chosen alternate amino acid with 10% probability, and the original input token (i.e., no change) with 10% probability.

#### B. One-hot encoding

The masked MSA is encoded as three matrices using one-hot encoding from three different views. Specifically, for the *j*-th position of the *i*-th sequence in the masked MSA, we encode it as three one-hot vectors, i.e., ***x***_*ij*_, ***y***_*ij*_, and ***z***_*ij*_, from the views of token type, row position, and column position, respectively.

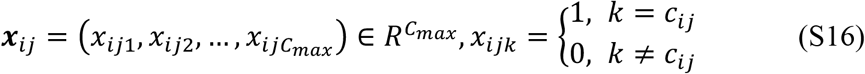

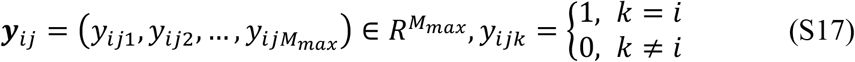

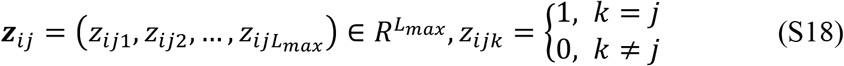

where *c*_*i*._ is the index of token type for the *j*-th position of the *i*-th sequence, *C*_*max*_ is the number of types of tokens, *L*_*max*_ and *M*_*max*_ are preset maximum values for sequence length and alignments, respectively. In this work, *C*_*max*_ = 28 and *L*_*max*_ = *M*_*max*_ = 1024, where 28 types of tokens include 20 common amino acids, 6 non-common amino acids (B, J, O, U, X and Z), 1 gap token, and 1 “masking” token.

According to Eqs. S16-S18, the masked MSA can be encoded as three matrices, i.e., ***X, Y*** and ***Z***, through one-hot encoding from the view of token type, row position, and column position, respectively, where 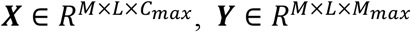 and 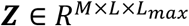, *M* is the number of alignments, and *L* is the length of individual sequence in the masked MSA.

#### C. Initial embedding

Each one-hot coding matrix is multiplied by a weight matrix to generate the corresponding embedding matrix:

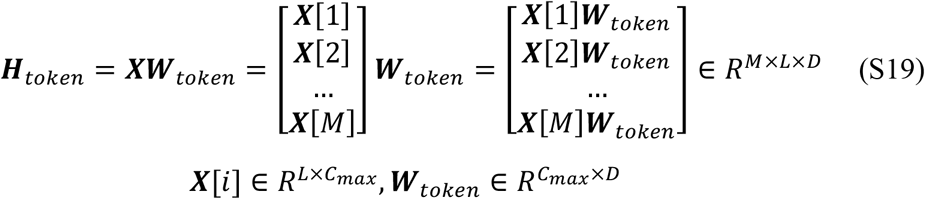

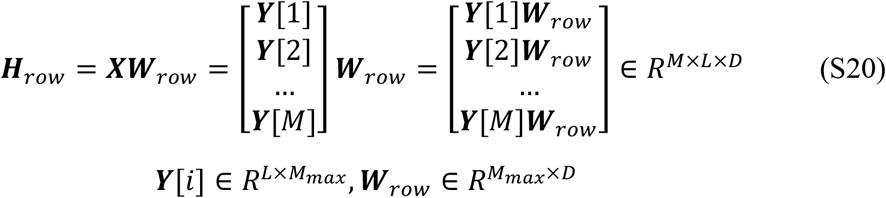

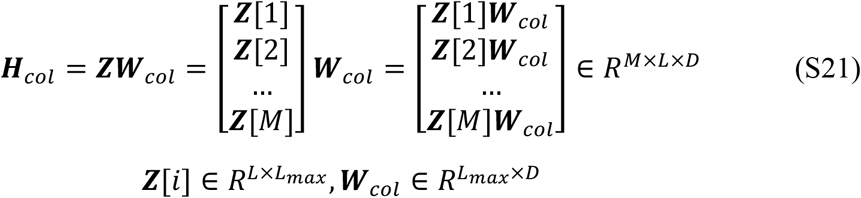

where ***X***[*i*], ***Y***[*i*] and ***Z***[*i*] are the one-hot coding matrices for the *i*-th sequence in the masked MSA from the view of token type, row position, and column position, respectively, ***H***_token_, ***H***_row_, and ***H***_col_ are token type-based, row position-based, and column position-based embedding matrices for the masked MSA, respectively, and *D* is the embedding dimension. In this work, *D* = 768.

Three embedding matrices are added as an initial embedding matrix ***H***_*init*_:

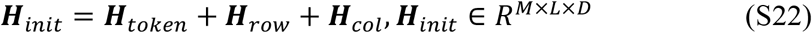

#### D. Batch normalization and dropout

The initial embedding matrix ***H***_*init*_ is fed to the batch normalization layer to generate the corresponding normalized matrix ***H***_1_:

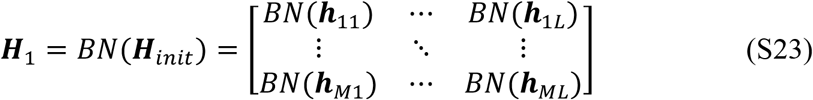

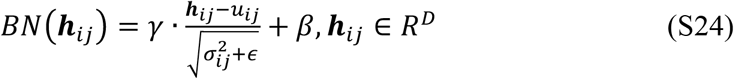

where ***h***_*ij*_ is the initial embedding vector for the *j*-th position of the *i*-th sequence in the masked MSA, *u*_*ij*_ and 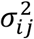 are mean and variance for ***h***_*ij*_, respectively, and *γ, β*, and *ϵ* are normalized factors.

The normalized matrix ***H***_1_ is fed to dropout layer:

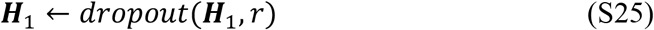

where *r* is the rate of neurons which are randomly dropped in each training step, indicating that the corresponding weight vectors will be not optimized.

#### E. Self-attention

The initial embedding matrix ***H***_1_ is fed to the self-attention network with *N* blocks, each of which consists of three sub-blocks. In this work, *N* = 12.

The first sub-block consists of a batch normalization layer, a row attention layer, a dropout layer, and a short connection, as follows.

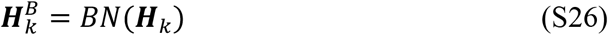

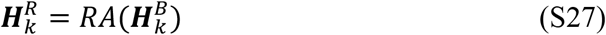

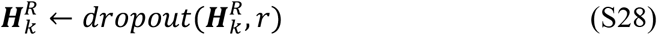

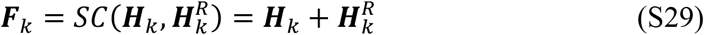

where ***H***_*k*_ and ***F***_*k*_ are the input and output matrices in the first sub-block of the *k*-th self-attention block, respectively, *BN*(∙) is the batch normalization function (see Eqs. S23-S24), *SC*(∙) is the short connection, and *RA*(∙) is the row attention layer (see Eqs. S38-S45), 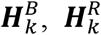.

The second sub-block consists of a batch normalization layer, a column attention layer, a dropout layer, and a short connection, as follows.

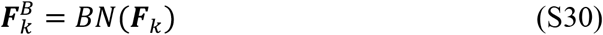

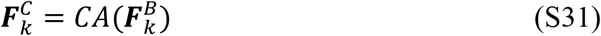

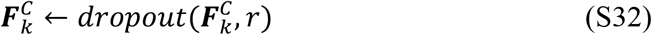

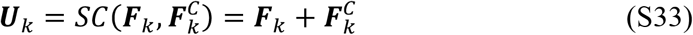

where ***F***_*k*_ and ***U***_*k*_ are the input and output matrices in the second sub-block of the *k*-th self-attention block, respectively, *CA*(∙) is the column attention layer (see Eqs. S46-S54), and 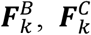.

The last sub-block consists of a batch normalization layer, a feed-forward network, a dropout layer, and a short connection, as follows.

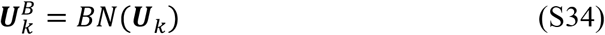

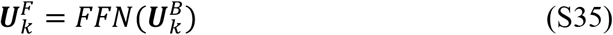

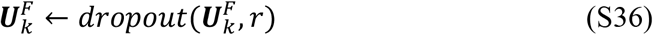

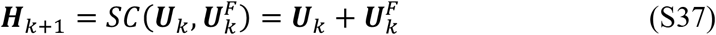

where ***U***_*k*_ and ***H***_*k*+1_ are the input and output matrices in the third sub-block of the *k*-th self-attention block, respectively, *FFN*(.) is the feed-forward network (see Eqs. S55-S60), and 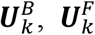.

#### (a) Row attention

Each row attention layer consists of *m* attention heads and a linear unit, where *m* = 12. In each attention head, the input matrix is multiplied by three weight matrices to generate the corresponding Query, Key, and Value matrices.

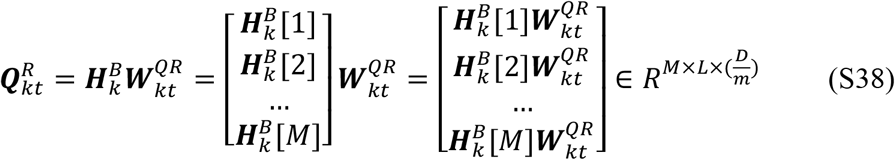

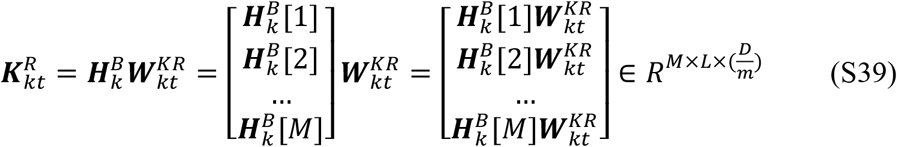

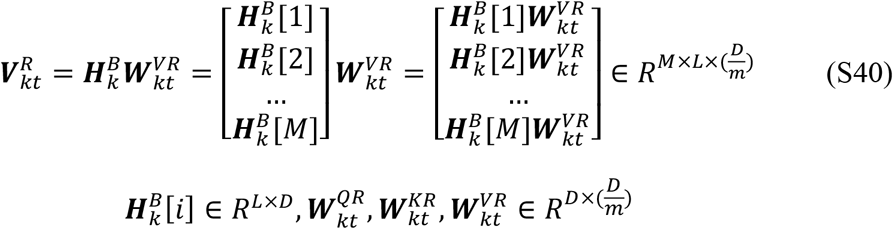

where 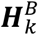 is the input matrix of row attention layer in the *k*-th self-attention block (See Eq. S27), 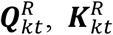, and 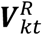 are Query, Key, and Value matrices in the *t*-th head of the row attention layer in the *k*-th block, respectively, 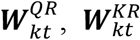, and 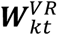 are corresponding weight metrices.

Then, the dot-product between 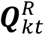 and 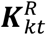 is performed and then normalized by SoftMax function to generate a row attention weight matrix:

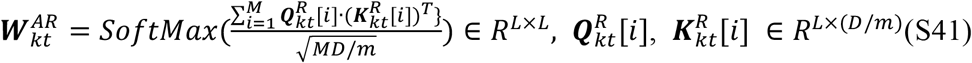

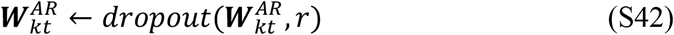

where 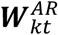 is the attention weight matrix in the *t*-th head of the row attention layer in the *k*-th block and measures the correlation for each pair of columns in the masked MSA.

Next, the row attention weight matrix 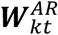 is multiplied by Value matrix 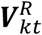 to generate the corresponding row attention matrix:

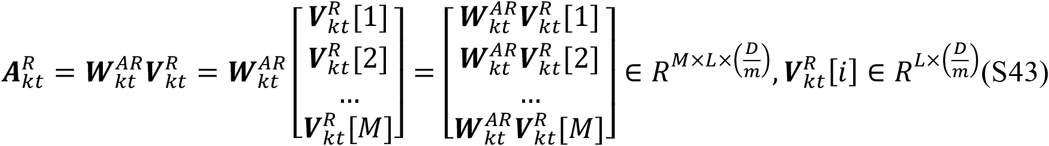

where 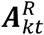 is the attention matrix in the *t*-th head of the row attention layer in the *k*-th block.

Finally, the outputs of all attention heads are concatenated as a new matrix, which is further fed to a linear unit:

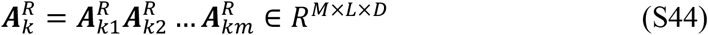

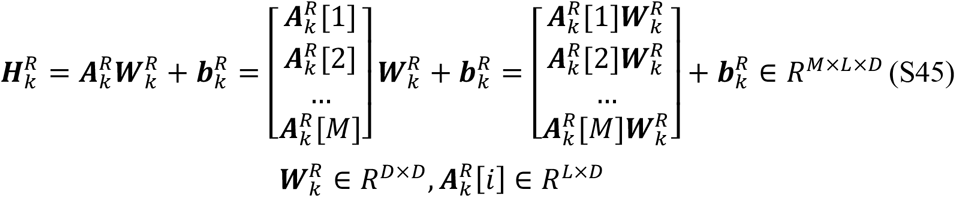

where 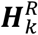 in the output matrix of row attention layer in the *k*-th attention block (See Eq. S27), and 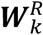 and 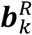 are weight matrix and bias in the linear unit, respectively.

#### (b) Column attention

Each column attention layer consists of *m* attention heads and a linear unit. In each attention head, the input matrix is multiplied by three weight matrices to generate the corresponding Query, Key, and Value matrices.

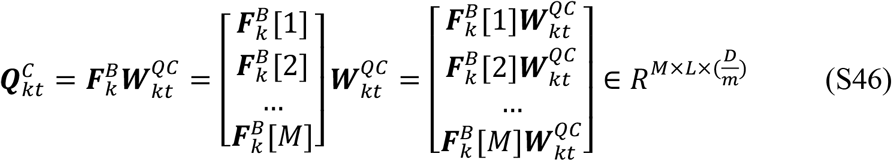

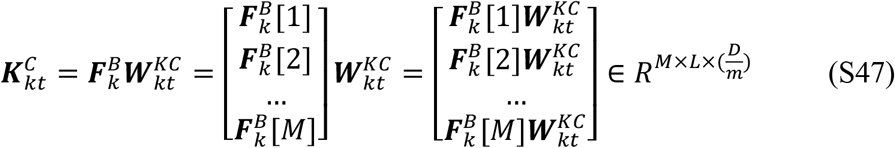

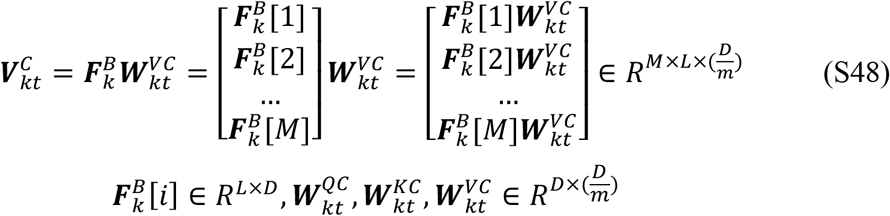

where 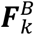 is the input matrix of column attention layer in the *k*-th self-attention block (see Eq. S31), 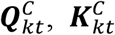, and 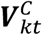 are Query, Key, and Value matrices in the *t*-th head of column attention layer in the *k*-th block, respectively, 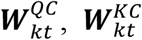, and 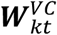 are corresponding weight metrices.

Then, the dot-product between 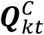 and 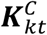 is performed and then normalized by SoftMax function to generate an attention weight matrix:

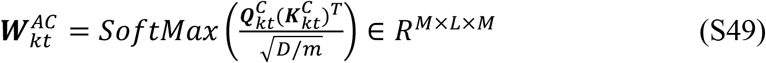

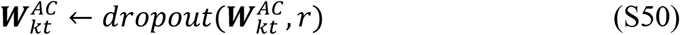

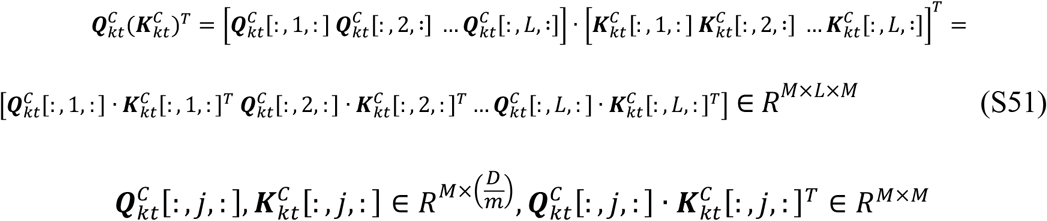

where 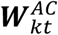 is the attention weight matrix in the *t*-th head of column attention layer in the *k*-th block, and 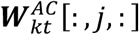 measures the correlation for each pair of alignments at the *j*-th position.

Next, the column attention weight matrix 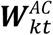 is multiplied by Value matrix 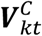 to generate the corresponding column attention matrix:

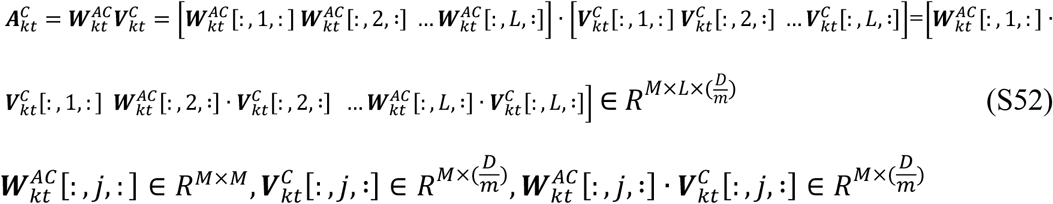

where 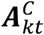 is the attention matrix in the *t*-th head of column attention layer in the *k*-th block.

Finally, the outputs of all attention heads are concatenated as a new matrix, which is further fed to a linear unit:

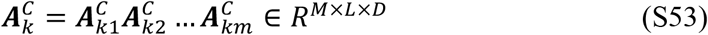

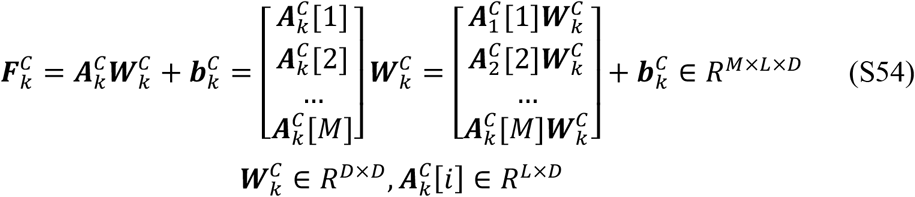

where 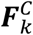 in the output matrix of column attention layer in the *k*-th attention block, (See Eq. S31), and 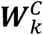 and 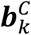 are weight matrix and bias in the linear unit, respectively.

#### (c) Feed-forward network

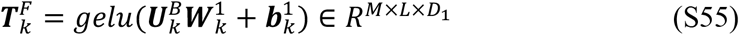

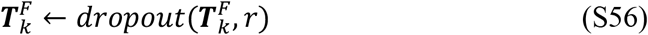

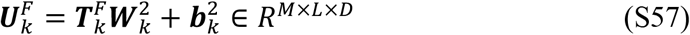

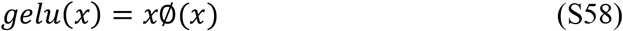

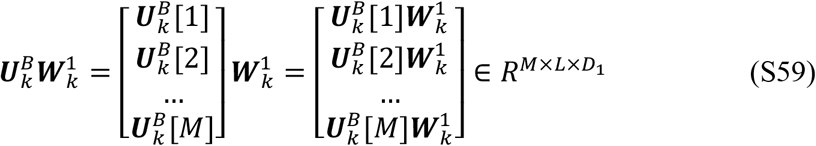

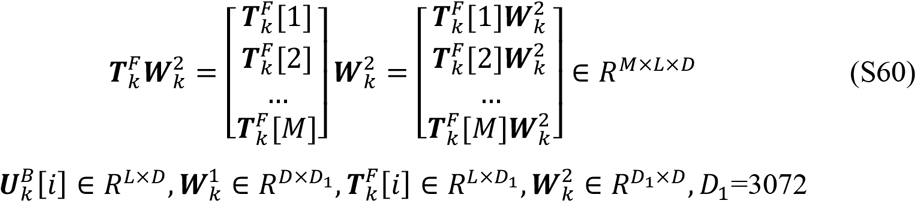

where 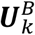 and 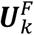 are the input and output matrices of feed-forward network in the *k* -th self-attention block, respectively, (see Eq. S35), 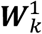 and 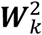 are weight matrices, 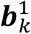 and 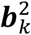 are bias, and ∅(*x*) is the integral of Gaussian Distribution for *x*.

#### F. Output layer

The output of the last self-attention block is fed to a fully connected layer with SoftMax function to generate a probability matrix:

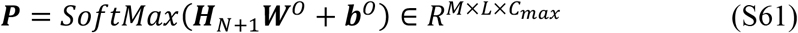

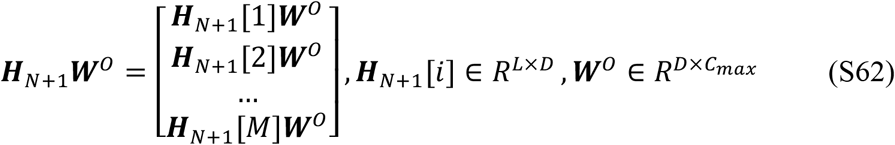

where ***H***_N+1_ is the outputted embedding matrix in the *N*-th self-attention block, ***W***° and ***b***° are weight matrix and bias, respectively, and the ***P***(*i, j, c*) indicates the probability that the *j*-th position of the *i*-th sequence in the masked MSA is predicted as the *c*-th type of amino acid.

#### G. Loss function

For an individual MSA, the loss function is designed as:

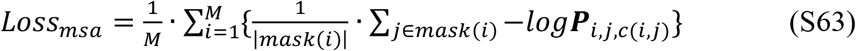

where *M* is the number of alignments, *mask*(*i*) is a set of masking position in the *i*-th sequence, |*mask*(*i*)| is the number of elements in *mask*(*i*), *c*(*i, j*) is the type index of amino acid for the *j*-th position in the *i*-th sequence before masking, and -*log****P***_*i,j*,c(*i,j*)_ is negative log likelihood of the true amino acid at the *j*-th position in the *i*-th sequence under condition of masking.

## Supporting Figures

**Figure S1.**
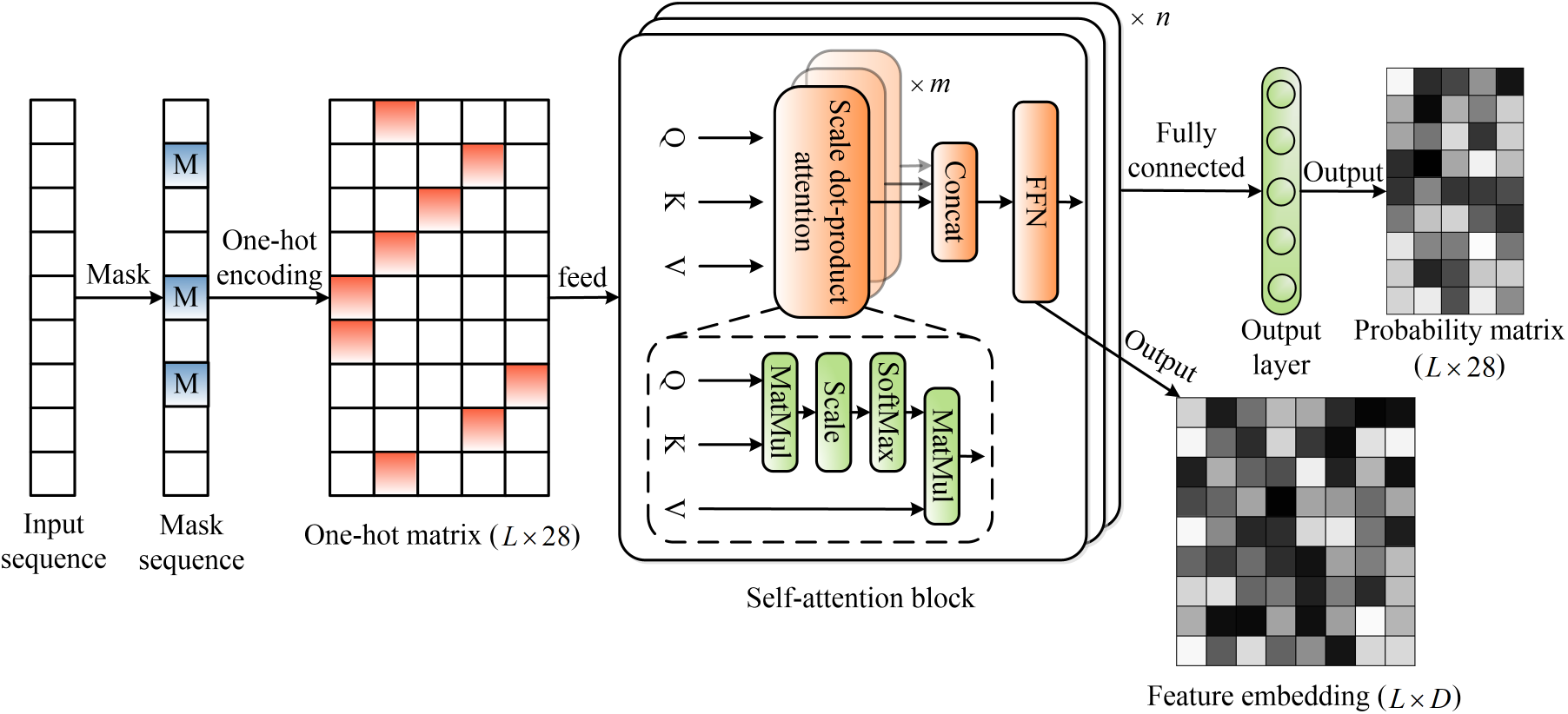
The workflow of ESM2 transformer.

**Figure S2.**
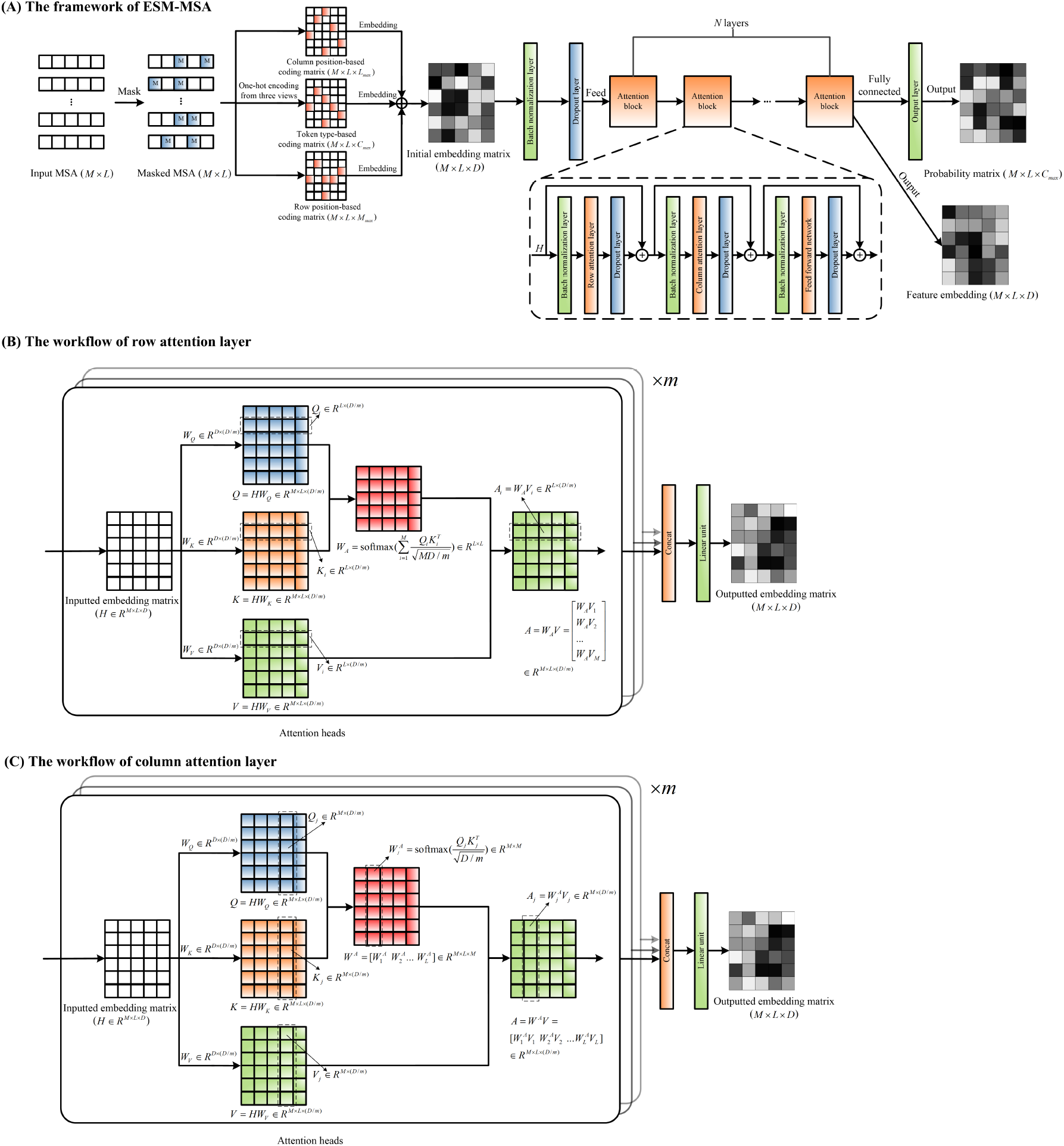
The workflow of ESM-MSA transformer.

**Figure S3.**
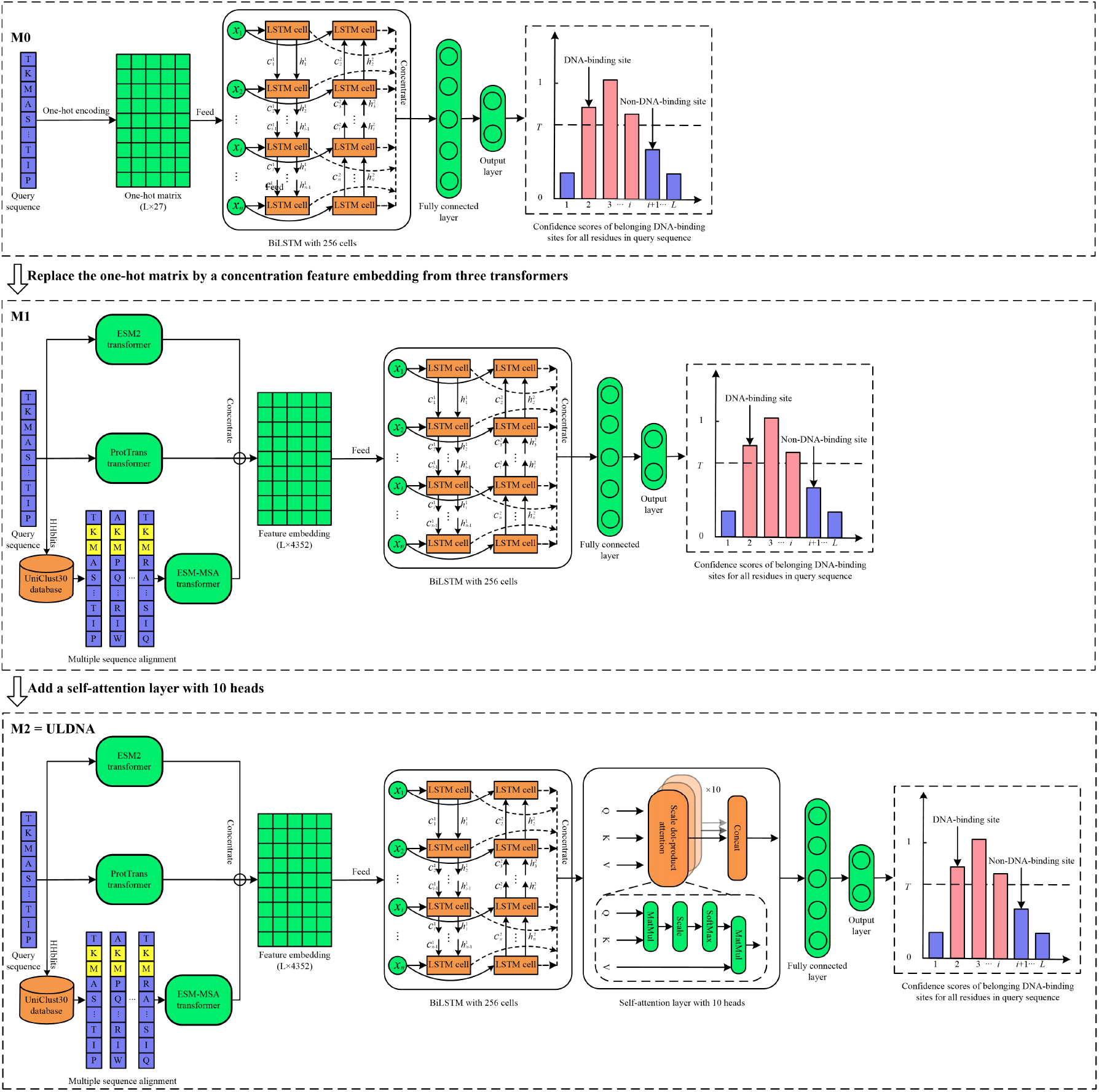
The architectures of three ablation models.

## Supporting Tables

**Table S1.** The performance of five DNA-binding site predictors on PDNA-543 over ten-fold cross-validation.

| Method | Sen | Spe | Acc | MCC | AUROC |
| --- | --- | --- | --- | --- | --- |
| TargetDNA ($Sen \approx Spe$)<sup>a</sup> | 0.770 | 0.771 | 0.770 | 0.304 | 0.845 |
| DNAPred ($Sen \approx Spe$)<sup>b</sup> | 0.771 | 0.785 | 0.784 | 0.318 | 0.861 |
| PredDBR ($Sen \approx Spe$)<sup>c</sup> | 0.776 | 0.774 | 0.774 | 0.358 | - |
| ULDNA ($Sen \approx Spe$) | **0.864** | **0.861** | **0.861** | **0.462** | **0.933** |
| TargetDNA ($Spe \approx 0.95$)<sup>a</sup> | 0.406 | 0.950 | 0.914 | 0.339 | 0.845 |
| DNAPred ($Spe \approx 0.95$)<sup>b</sup> | 0.449 | 0.950 | 0.917 | 0.373 | 0.861 |
| PredDBR ($Spe \approx 0.95$)<sup>c</sup> | 0.454 | **0.955** | 0.914 | 0.415 | - |
| Guan's method ($Spe \approx 0.95$)<sup>d</sup> | 0.452 | 0.954 | 0.928 | 0.352 | - |
| ULDNA ($Spe \approx 0.95$) | **0.668** | 0.950 | **0.931** | **0.534** | **0.933** |

<sup>a</sup> Results excerpted from TargetDNA [4]; <sup>b</sup> Results excerpted from DNAPred [5]; <sup>c</sup> Results excerpted from PredDBR [6]; <sup>d</sup> Results excerpted from Guan et al [7]. " $Sen \approx Spe$ " means the threshold that makes $Sen \approx Spe$ ; " $Spe \approx 0.95$ " means the threshold that makes $Spe \approx 0.95$ . '-' means the value is not given. Bold fonts highlight the best performer in each evaluation metric.

**Table S2.** The performance of five DNA-binding site predictors on PDNA-335 over five-fold cross-validation.

| Method | Sen | Spe | Acc | MCC | AUROC |
| --- | --- | --- | --- | --- | --- |
| EC-RUS<sup>a</sup> | 0.487 | 0.951 | **0.926** | 0.378 | 0.852 |
| TargetS<sup>b</sup> | 0.417 | 0.945 | 0.899 | 0.362 | 0.824 |
| DNAPred<sup>c</sup> | 0.543 | 0.917 | 0.886 | 0.390 | 0.856 |
| PredDBR<sup>d</sup> | 0.426 | **0.953** | 0.910 | 0.390 | - |
| ULDNA | **0.676** | 0.948 | 0.925 | **0.565** | **0.940** |

<sup>a</sup> Results excerpted from EC-RUS [8]; <sup>b</sup> Results excerpted from TargetS [9]; <sup>c</sup> Results excerpted from DNAPred [5]; <sup>d</sup> Results excerpted from PredDBR [6]. '-' means the value is not given. Bold fonts highlight the best performer in each evaluation metric.

**Table S3.**
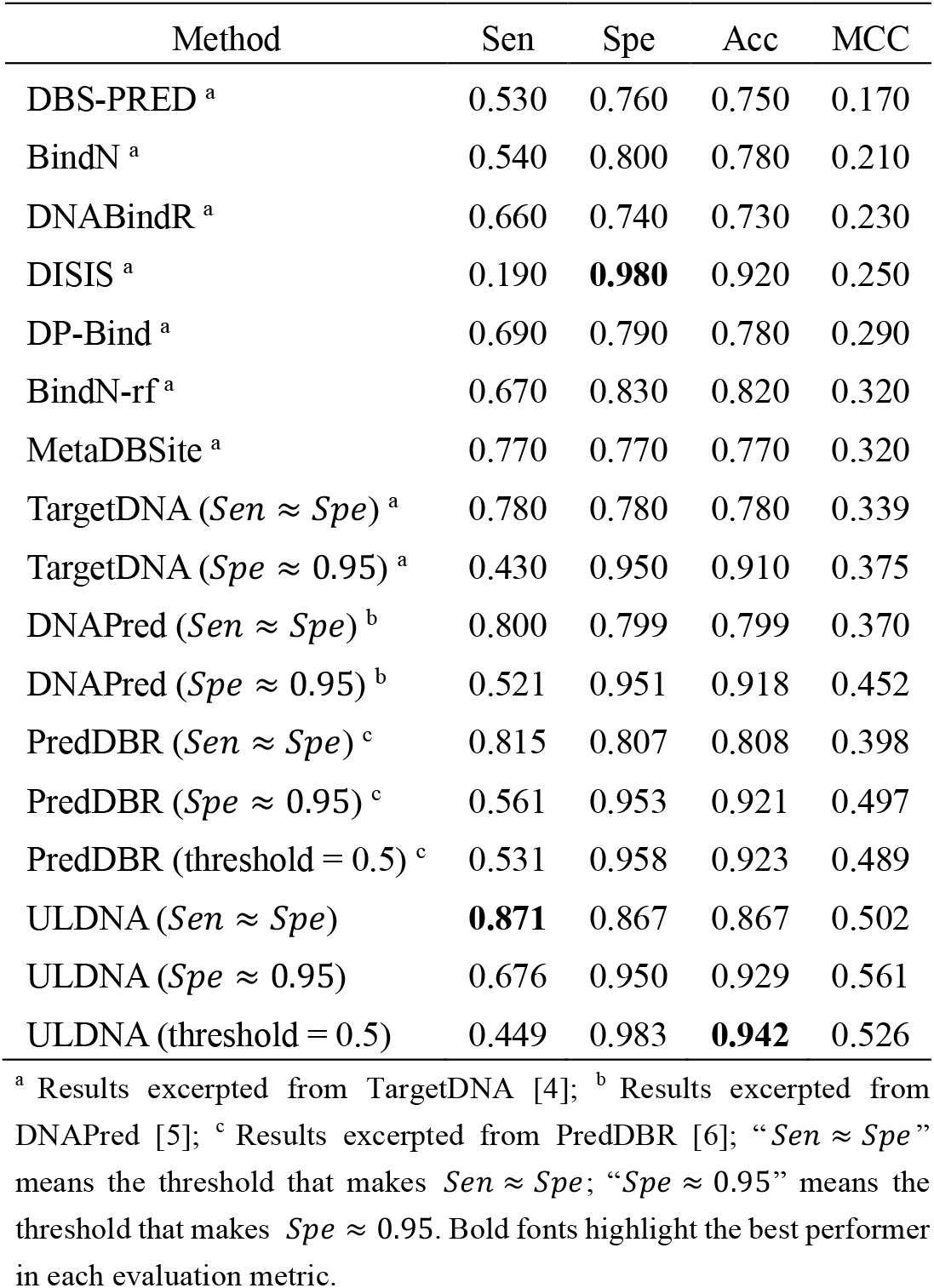
The performance of eleven DNA-binding site predictors on PDNA-316 over ten-fold cross-validation.

| Method | Sen | Spe | Acc | MCC |
| --- | --- | --- | --- | --- |
| DBS-PRED <sup>a</sup> | 0.530 | 0.760 | 0.750 | 0.170 |
| BindN <sup>a</sup> | 0.540 | 0.800 | 0.780 | 0.210 |
| DNABindR <sup>a</sup> | 0.660 | 0.740 | 0.730 | 0.230 |
| DISIS <sup>a</sup> | 0.190 | **0.980** | 0.920 | 0.250 |
| DP-Bind <sup>a</sup> | 0.690 | 0.790 | 0.780 | 0.290 |
| BindN-rf <sup>a</sup> | 0.670 | 0.830 | 0.820 | 0.320 |
| MetaDBSite <sup>a</sup> | 0.770 | 0.770 | 0.770 | 0.320 |
| TargetDNA (*Sen* ≈ *Spe*) <sup>a</sup> | 0.780 | 0.780 | 0.780 | 0.339 |
| TargetDNA (*Spe* ≈ 0.95) <sup>a</sup> | 0.430 | 0.950 | 0.910 | 0.375 |
| DNAPred (*Sen* ≈ *Spe*) <sup>b</sup> | 0.800 | 0.799 | 0.799 | 0.370 |
| DNAPred (*Spe* ≈ 0.95) <sup>b</sup> | 0.521 | 0.951 | 0.918 | 0.452 |
| PredDBR (*Sen* ≈ *Spe*) <sup>c</sup> | 0.815 | 0.807 | 0.808 | 0.398 |
| PredDBR (*Spe* ≈ 0.95) <sup>c</sup> | 0.561 | 0.953 | 0.921 | 0.497 |
| PredDBR (threshold = 0.5) <sup>c</sup> | 0.531 | 0.958 | 0.923 | 0.489 |
| ULDNA (*Sen* ≈ *Spe*) | **0.871** | 0.867 | 0.867 | 0.502 |
| ULDNA (*Spe* ≈ 0.95) | 0.676 | 0.950 | 0.929 | 0.561 |
| ULDNA (threshold = 0.5) | 0.449 | 0.983 | **0.942** | 0.526 |

<sup>a</sup> Results excerpted from TargetDNA [4]; <sup>b</sup> Results excerpted from DNAPred [5]; <sup>c</sup> Results excerpted from PredDBR [6]; “*Sen* ≈ *Spe*” means the threshold that makes *Sen* ≈ *Spe*; “*Spe* ≈ 0.95” means the threshold that makes *Spe* ≈ 0.95. Bold fonts highlight the best performer in each evaluation metric.

**Table S4.**
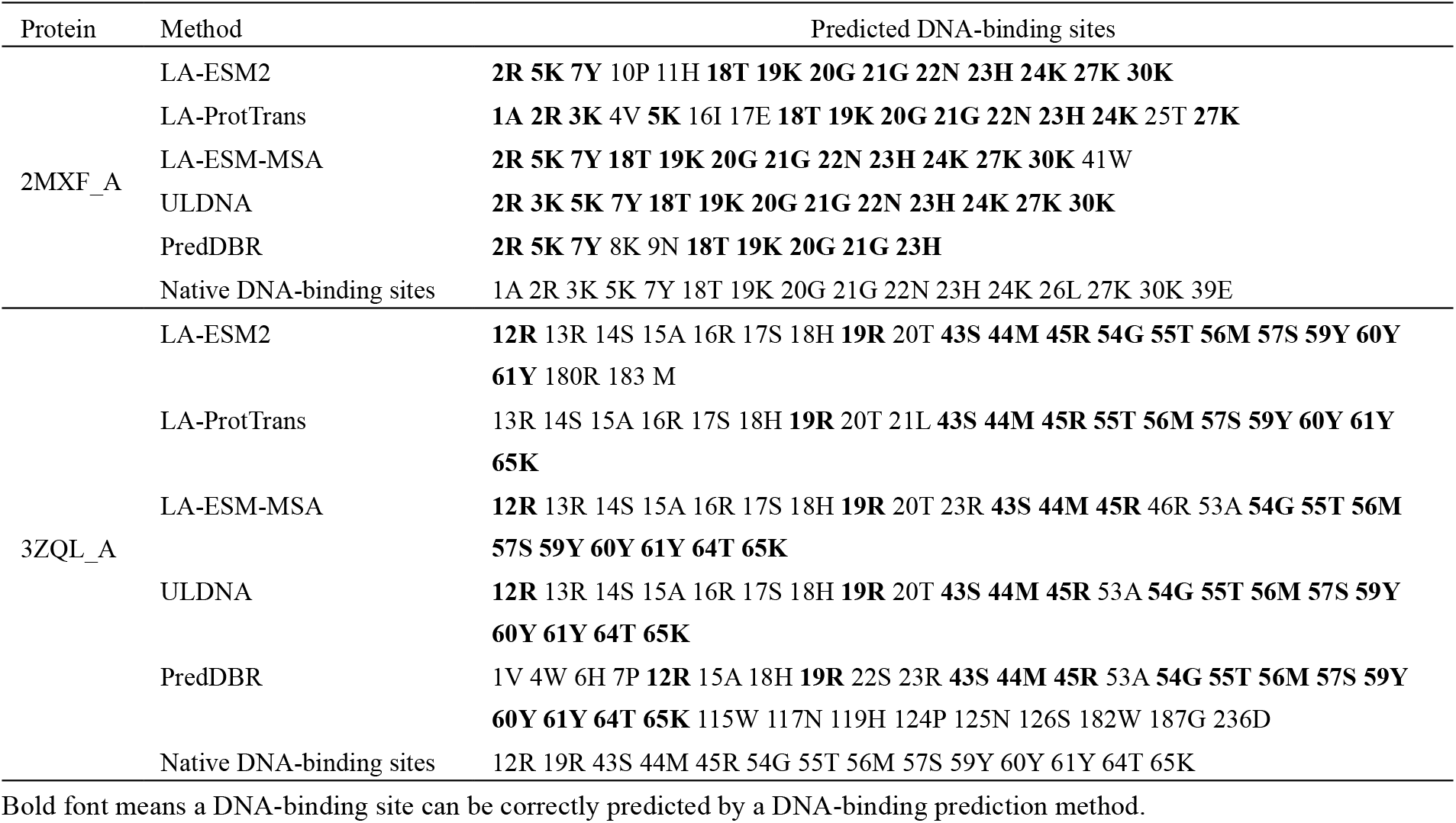
The predicted and native DNA-binding sites of two representative proteins for five DNA-binding prediction methods

